# MTaxi : A comparative tool for taxon identification of ultra low coverage ancient genomes

**DOI:** 10.1101/2022.06.06.491147

**Authors:** Gözde Atağ, Kıvılcım Başak Vural, Damla Kaptan, Mustafa Özkan, Dilek Koptekin, Ekin Sağlıcan, Sevcan Doğramacı, Mevlüt Köz, Ardan Yılmaz, Arda Söylev, İnci Togan, Mehmet Somel, Füsun Özer

**Affiliations:** Department of Biological Sciences, Middle East Technical University, Ankara, Turkey; Department of Health Informatics, Graduate School of Informatics, Middle East Technical University, Ankara, Turkey; Department of Anthropology, Hacettepe University, Ankara, Turkey; Department of Computer Engineering, Konya Food and Agriculture University, Konya, Turkey; Department of Molecular Biology and Genetics, Konya Food and Agriculture University, Konya, Turkey; Department of Computer Engineering, Middle East Technical University, Ankara, Turkey

## Abstract

A major challenge in zooarchaeology is to morphologically distinguish closely related species’ remains, especially using small bone fragments. Shotgun sequencing aDNA from archeological remains and comparative alignment to the candidate species’ reference genomes will only apply when reference nuclear genomes of comparable quality are available, and may still fail when coverages are low. Here, we propose an alternative method, MTaxi, that uses highly accessible mitochondrial DNA (mtDNA) to distinguish between pairs of closely related species from ancient DNA sequences. MTaxi utilises mtDNA transversion-type substitutions between pairs of candidate species, assigns reads to either species, and performs a binomial test to determine the sample taxon. We tested MTaxi on sheep/goat and horse/donkey data, between which zooarchaeological classification can be challenging in ways that epitomise our case. The method performed efficiently on simulated ancient genomes down to 0.5x mitochondrial coverage for both sheep/goat and horse/donkey, with no false positives. Trials on n=18 ancient sheep/goat samples and n=10 horse/donkey samples of known species identity with mtDNA coverages 0.1x - 12x also yielded 100% accuracy. Overall, MTaxi provides a straightforward approach to classify closely related species that are compelling to distinguish through zooarchaeological methods using low coverage aDNA data, especially when similar quality reference genomes are unavailable. MTaxi is freely available at https://github.com/goztag/MTaxi.

## Introduction

Archaeological faunal remains have been widely used to address various questions in biology and social sciences. The scope of these range from the demographic history of wild populations, which can inform about ecological dynamics and conservation biology, to animal management and breeding practices, providing insights into the subsistence strategies and lifeways of prehistoric human societies that exploited animals (1-6). A key step here is the accurate taxonomic identification of animal remains. However, distinguishing morphologically similar species in zooarchaeological material is a prevailing challenge, constrained by the high level of similarity between skeletal elements, the fragmented state of excavated specimens (possibly with missing fragments), and the absence of morphological markers in subadults (7,8). The need for an effective approach to identify species’ remains accurately has thus led to the development of several alternative methods, including isotope analyses, protein fingerprinting, and ancient DNA (aDNA) analyses (9-14).

The majority of non-human aDNA data today is produced using shotgun DNA sequencing on Illumina platforms (15). Beyond species identification, such data from well-preserved zooarchaeological samples can yield a wealth of information to study demographic and evolutionary history. However, relatively old (e.g. >1000 years old) zooarchaeological samples from regions with humid, temperate or warmer environments are mostly poorly preserved (16). Cooking and other forms of heat treatment before human consumption may additionally degrade organic material (17). In such poorly preserved samples, the proportion of endogenous DNA among the total DNA read pool will be low, down to 1% or even lower (18). Accordingly, most experiments can produce only low amounts of DNA sequence data, if any, from zooarchaeological samples from temperate regions within reasonable budgets; such genomic data frequently remain at genome-wide depths of coverage <0.1x per sample (19, 20).

Theoretically, even 0.1x coverage genome data could allow accurate taxonomic identification by comparative alignment, i.e. mapping reads to the reference genomes of alternative candidate species, such as sheep versus goat, or horse versus donkey, and comparing coverages or mismatch rates. However, this only applies to situations where both species have assembled nuclear reference genomes (e.g. no such reference is available for the donkey). Even in cases where nuclear genomes from both species are available (e.g. sheep and goat), the limited amount of shotgun sequencing data available from poorly preserved samples, quality differences between the reference genomes, the highly fragmented nature of aDNA hence short read lengths, and postmortem damage can introduce high levels of uncertainty in the alignment process. The problem is further exacerbated when the sequence similarity between candidate reference genomes is high or when there exists strong differences in the genome assemblage qualities.

These call for new approaches for species identification with aDNA data. For instance, the Zonkey (21) pipeline was developed for distinguishing horse, donkey and their hybrids by using nuclear aDNA variants with a clustering approach, but is only applicable for equid taxa from which there exists large datasets of genetic variation. Here we present a broadly applicable method, MTaxi, designed for distinguishing pairs of any closely related species using low amounts of shotgun aDNA sequencing data, whenever mitochondrial DNA (mtDNA) reference sequences are available. Our method focuses on mtDNA owing to its short size, haploid nature, having a lower rate of decay than nuclear DNA (16), and having multi-copies per cell, which increases its availability relative to autosomes (22), facilitating analyses. For example, across n=310 shotgun sequenced ancient DNA libraries from human, sheep, goat, horse and donkey generated by our group, each of which contained ≥0.01 endogenous DNA, the average ratio of mitochondrial DNA to nuclear DNA coverage was 87:1 (data not shown). The greater number of informative sites due to the high mitochondrial mutation rate is an additional advantage for taxon identification of closely related species (23). Finally, the availability of mitochondrial reference sequences for a larger number of taxa (compared to a limited number of high quality reference genomes) allows our approach to be applied to a wider number of species, including extinct lineages. For instance, as of December 24 2021, the genome resources database from NCBI includes only 175 nuclear genomes for mammals (24), compared to 1453 mitochondrial genomes (25), an 8-fold difference.

To exemplify the use of MTaxi we chose the case of sheep (Ovis) versus goat (Capra), two closely related species belonging to the same subfamily Caprinae. The aforementioned constraints on morphological identification causes a large proportion of sheep and goat remains to be only identified at the subfamily level as Caprinae (8), and ambiguity which can significantly constrain zooarchaeological analyses, especially in the study of animal husbandry. Here we first estimate MTaxi’s accuracy using 1200 ancient mitogenome simulations with six different coverages from both sheep and goat. We then test its performance with n=9 ancient sheep samples (19,20) and n=3 goat samples (26). We further test MTaxi on the horse and donkey, a pair which also suffer from difficulties in morphological differentiation, using n=3 ancient horse, n=2 modern domestic horse, and n=5 modern domestic donkey samples adopted from the literature (27-31).

## Materials and methods

### Overview of the method

MTaxi makes use of the mismatch positions, i.e. putative substitutions, between two alternative candidate taxa, such as sheep and goat. We call these “target sites” and are obtained from pairwise alignment of mitochondrial reference genomes. Each read harbouring the target sites is classified according to the genotype compositions, and we identify the taxon using a binomial test for the read proportion of the sample (Figure 1).

**Figure 1.**
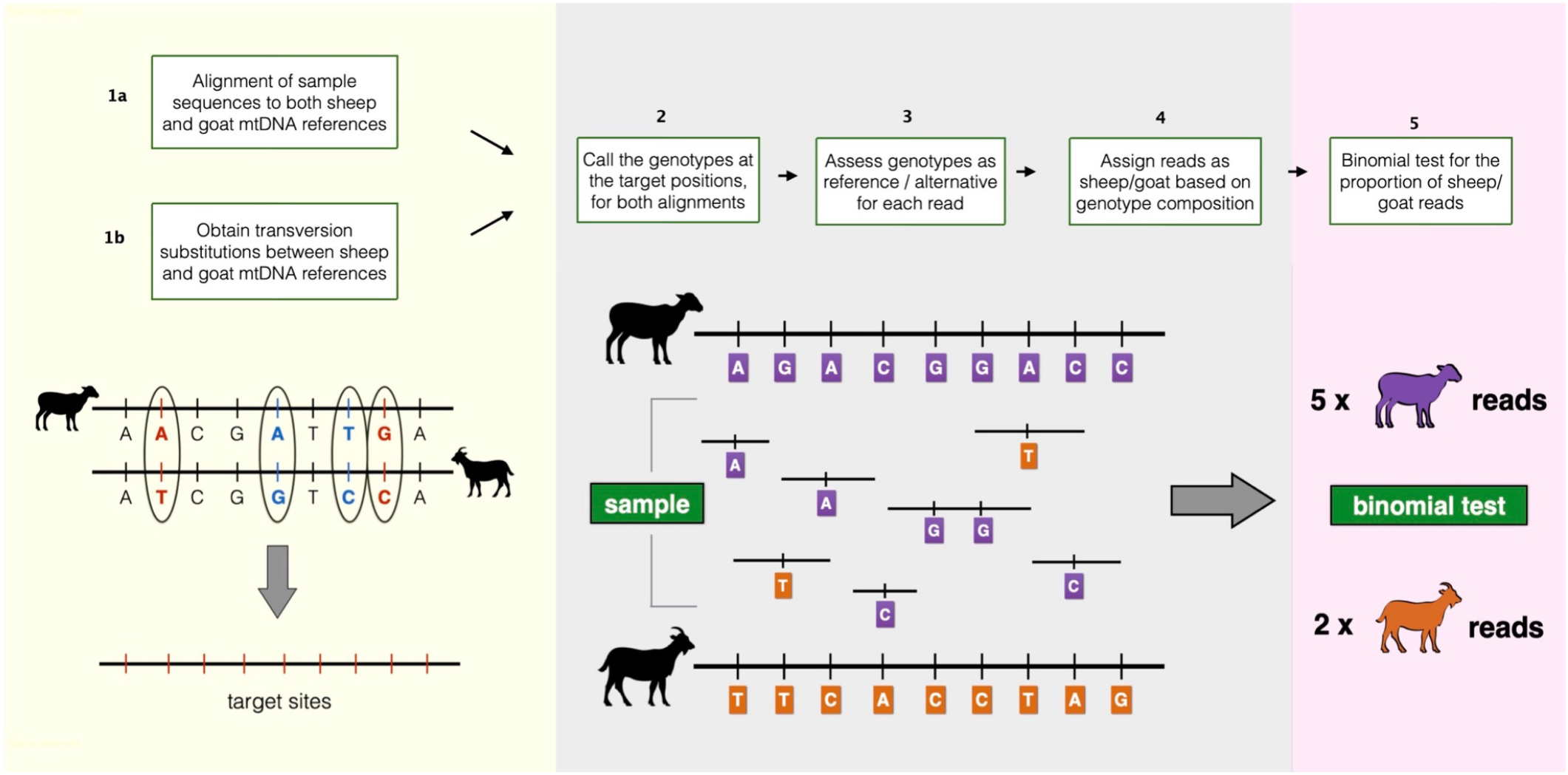
Overview of the MTaxi pipeline. Flowchart and representations of the steps to determine the sample taxon. Here sheep and goat stand for the candidate species pair, but MTaxi can be applied to any pair of species where mitochondrial DNA reference sequences are available. Target sites represent mismatches (candidate substitutions) between the reference genomes, restricted to transversions. Reads are assigned to either taxon based on target sites. Reads may be assigned to the wrong taxon due to homoplastic mutations, technical error, or incomplete lineage sorting.

### Target sites

The method involves compiling a list of mtDNA target sites, representing likely substitutions between the species. To generate this list for sheep and goat, we first generated a pairwise alignment between sheep (Oar_v3.1) and goat (ARS1) mtDNA reference genomes via the R package *Biostrings* v.2.65.0 (32) using default parameters, which yielded n=1699 single nucleotide substitutions. We then restricted these to transversions to avoid (a) confounding effects due to postmortem damage-induced transitions in ancient DNA, and (b) homoplasies that could arise by high-frequency transitions (33). This yielded a set of n=197 transversion substitutions, which we refer to as the target site list 1.

We also created a subset of this, that we call target sites type 2, by removing polymorphisms in either species, which we reasoned might increase power by avoiding ambiguities. For this, we obtained a list of polymorphic sites using the software *snp-sites* v.2.4.1 (34) from a data set assembled by Shi and colleagues (35), which contains pairwise alignments (each sequence aligned to the reference genome) for mtDNA sequences belonging to n=47 domestic sheep and n=35 domestic goats. In this dataset, we identified n=57 and n=40 polymorphic single nucleotide positions overlapping with the n=197 target sites in domestic sheep and goat, respectively. After eliminating these polymorphisms we were left with n=120 positions, which we refer to as the target site list 2.

We applied the same procedures to horse and donkey by using mtDNA references NC_001640.1 and NC_001788.1. This resulted in n=1264 substitution sites, and restricting these to transversions yielded n=117 positions (Figure 2). The positions are concentrated around the D-loop, but are also represented across the mitochondrial genome following similar patterns between the two species.

**Figure 2.**
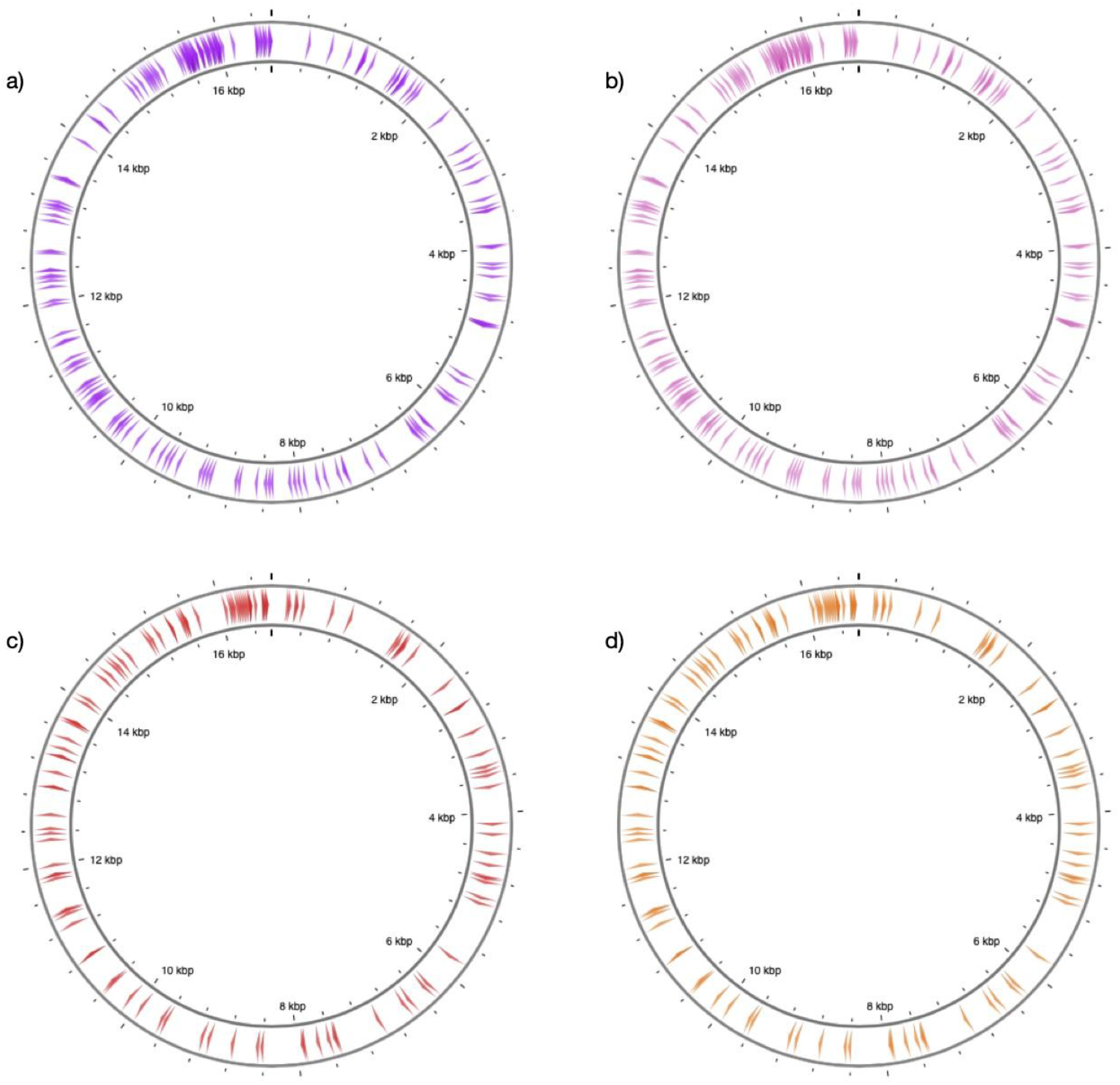
Distribution of target sites along reference mitochondrial genomes. The figure shows the position of target sites along (a) sheep and (b) goat (c) horse (d) donkey reference mitochondrial genomes. The sites represent transversion-type substitutions (n=197 for sheep and goat, and n=117 for horse and donkey). The figure was generated through CGview (36).

### Alignment and genotyping

#### Ancient DNA data processing

*AdapterRemoval* v.2.3.1 (37) was used for trimming residual adapter sequences and merging paired-end sequencing reads with an overlap of at least 11 bp between the pairs. Whole genome sheep and goat data (FASTQ files) were aligned to both sheep (Oar_v3.1) and goat (ARS1) reference genomes using *BWA aln* v.0.7.15 (38) with parameters: “-n 0.01 -o 2” and by disabling the seed with “-l 16500”. Mitochondrial goat data (Gilat10, Shiqmim9, Kov27, Uiv17) were aligned to both sheep (NC_001941.1) and goat (NC_005044.2) mitochondrial references with the same parameters as well. To prevent the influence of the PCR duplicates, reads with identical start and end positions were removed using “FilterUniqSAMCons_cc.py” (39). After the removal of PCR duplicates, reads with mapping quality scores (MAPQ) lower than 30 were filtered out using *samtools* (v.1.9) (40). The reads mapping to the reference genome with >10% mismatches and having a length <35 bp were filtered out. Damage patterns which are characteristics of aDNA were estimated using *PMDtools* (41) “--deamination” parameter. The reads aligned to mtDNA were extracted from whole genome alignments using *samtools* (v.1.9) (40). For each alignment, we called the genotypes at our sites of interest using *pysam* v.0.16.0.1 (https://github.com/pysam-developers/pysam), which also runs *samtools* (v.1.9) (40); genotyping was performed with parameters “-B” and “-A”. As default, MTaxi uses both the reads that aligned only to one of the species’ references and the ones that aligned to both species’ references in the analysis (which we refer to as “all reads” below). Additionally, we included an option (“shared reads”), by which the reads are restricted to those that are aligned to both species’ references; this is a conservative approach that could eliminate the possible effects of quality differences between the two reference genomes. Using *pybedtools* v.0.8.1 (42,43), we obtained the reads overlapping with the target sites. We note that aligning reads to both nuclear and mitochondrial genomes is superior to alignment only to the mitochondrial genome, because the latter can cause misalignment of nuclear mitochondrial DNA sequences (NUMTs) to the mitochondrial genome.

The alignment and genotyping procedures for whole genome ancient horse data were applied in the same way as described for sheep and goat data. However, for the alignment, due to lack of a nuclear reference genome for the donkey, equid reads were mapped only to mtDNA references of the two species.

For the comparison of whole genome mapping frequencies, the total number of bases aligned to both reference genomes were calculated using *samtools stats* (40), and we calculated the number of mismatches for each alignment.

#### Modern DNA data processing

After removing residual adapter sequences using *AdapterRemoval* v.2.3.1 (37), we mapped the whole genome data of modern horse and donkey at pair-ended mode to both horse (NC_001640.1) and donkey (NC_001788) mitochondrial reference genomes using *BWA mem* (version 0.7.15) (38) module with the parameter ‘-M’, and sorted the output using *samtools* (v.1.9) *sort* (40). Duplicates were removed using *Picard MarkDuplicates* (http://broadinstitute.github.io/picard/). Reads with mapping quality scores lower than 20 were filtered out using *samtools* (v.1.9) (40). Libraries from the same individual were merged using *samtools* (v.1.9) *merge* (40), and then the same filtering and genotyping procedures described in Ancient DNA data processing section process were applied on all modern equid data.

### Taxon assignment

Each read can carry either reference or alternative genotypes at its target sites. MTaxi uses this genotype data to assign reads to either taxon, species 1 (SP1) or species 2 (SP2). This is done by first calculating the frequency of alternative alleles per read. If an SP1 read was aligned to the SP1 genome, we expect no alternative alleles at target sites, and if aligned to the SP2 genome, we expect all alternative alleles. MTaxi retains reads with an alternative allele frequency of either 1 or 0, thus excluding reads with inconsistent genotypes (i.e. alternative and reference alleles mixed) at target sites. Such inconsistent variants could represent PCR or sequencing errors, convergent mutations, or incomplete lineage sorting. Having thereby assigned reads as SP1 or SP2, MTaxi uses the proportion of these two classes of reads to determine the sample taxon using a two-tailed binomial test with the null hypothesis of p=0.5.

### Ancient mitogenome simulations

We simulated 1200 ancient sheep and goat mitochondrial genomes (100 sheep and 100 goats for each coverage) at six different coverages (0.5x, 1x, 2x, 3x, 4x, 5x) using *gargammel* (44), and tested the accuracy of the method. The sequencing error was set to ∼1% using the parameters “qs -10 qs2 -10”. The simulations for horse and donkey (100 horse and 100 donkey for each coverage) were run with the same parameters above, again at six different coverages (0.5x,1x,2x,3x,4x,5x). The same alignment and genotyping procedures described in Ancient DNA data processing section were applied to the simulated data, except that they were mapped only to the mtDNA references of the species.

### Ancient and modern samples

We used published FASTQ files to study the performance of MTaxi on real data. For sheep, we used FASTQ files of n=5 ancient sheep individuals (TEP03, TEP62, TEP83, ULU26, ULU31) from Yurtman et al. (19) downloaded from the European Nucleotide Archive (ENA) database (Table 1), and n=4 (OBI013,OBI014, OBI017, OBI018) ancient sheep individuals from Taylor et al. (20) downloaded from ENA (Table 1). All data had been produced with Illumina sequencing using either whole genome shotgun sequencing or using SNP capture followed by sequencing. For goat, we used n=5 (Acem1, AP45, Azer3, Direkli1, Direkli6) ancient whole genome FASTQ files, and n=4 (Gilat10, Shiqmim9, Kov27, Uiv17) ancient mitochondrial capture FASTQ files, produced by shotgun Illumina sequencing and mtDNA capture-sequencing, and published by Daly et al. (26), downloaded from ENA (Table 1). For equids, we used n=3 ancient (27) and n=2 modern domestic horses (28,29) and n=5 modern domestic donkey (30,31) FASTQ files, downloaded from ENA (Table 1). We randomly downsampled the equid files to mtDNA coverages ranging from ∼0.3x to ∼4x using *samtools view* with the option “-s” (40).

**Table 1.** Genome data used in the study. The table lists ancient and modern-day genomes downloaded from European Nucleotide Archive (ENA), with the study accession IDs and sample aliases.

| Sample Aliases | Species | Age | Study Accession | Publication |
| --- | --- | --- | --- | --- |
| TEP03, TEP62, TEP83, ULU26, ULU31 | Sheep | Ancient | PRJEB36540 | (19) |
| OBI013,OBI014, OBI017, OBI018 | Sheep | Ancient | PRJEB41594 | (20) |
| Acem1, AP45, Azer3, Direkli1, Direkli6, Gilat10, Shiqmim9, Kov27, Uiv17 | Goat | Ancient | PRJEB26011 | (26) |
| Au6, Et1, Ke14, Sp5 | Donkey | Modern | PRJNA431818 | (31) |
| Willy | Donkey | Modern | PRJEB24845 | (30) |
| FM1798 | Horse | Modern | PRJEB10098 | (29) |
| Twilight | Horse | Modern | PRJNA205517 | (28) |
| VHR031, VHR102, CdY2 | Horse | Ancient | PRJEB31613 | (27) |

## Results

### Application to simulated ancient mitogenomes

We first studied the performance of MTaxi using ancient-like mtDNA read data simulations. We produced n=1200 mtDNA read datasets at varying coverage, n=600 for sheep and n=600 for goat (Materials and Methods). Using n=197 transversion substitutions (target sites type 1), MTaxi assigned BAM files to their respective taxa with 100% precision (i.e. no false positives) across all mtDNA coverages from 0.5x-5x using the default (“all reads”) approach (Figure 3a,b). All simulated data had a recall (i.e. true positive rate) of 100% and no false positives, even at mtDNA coverages ≥0.5x (Figure 3a).

**Figure 3.**
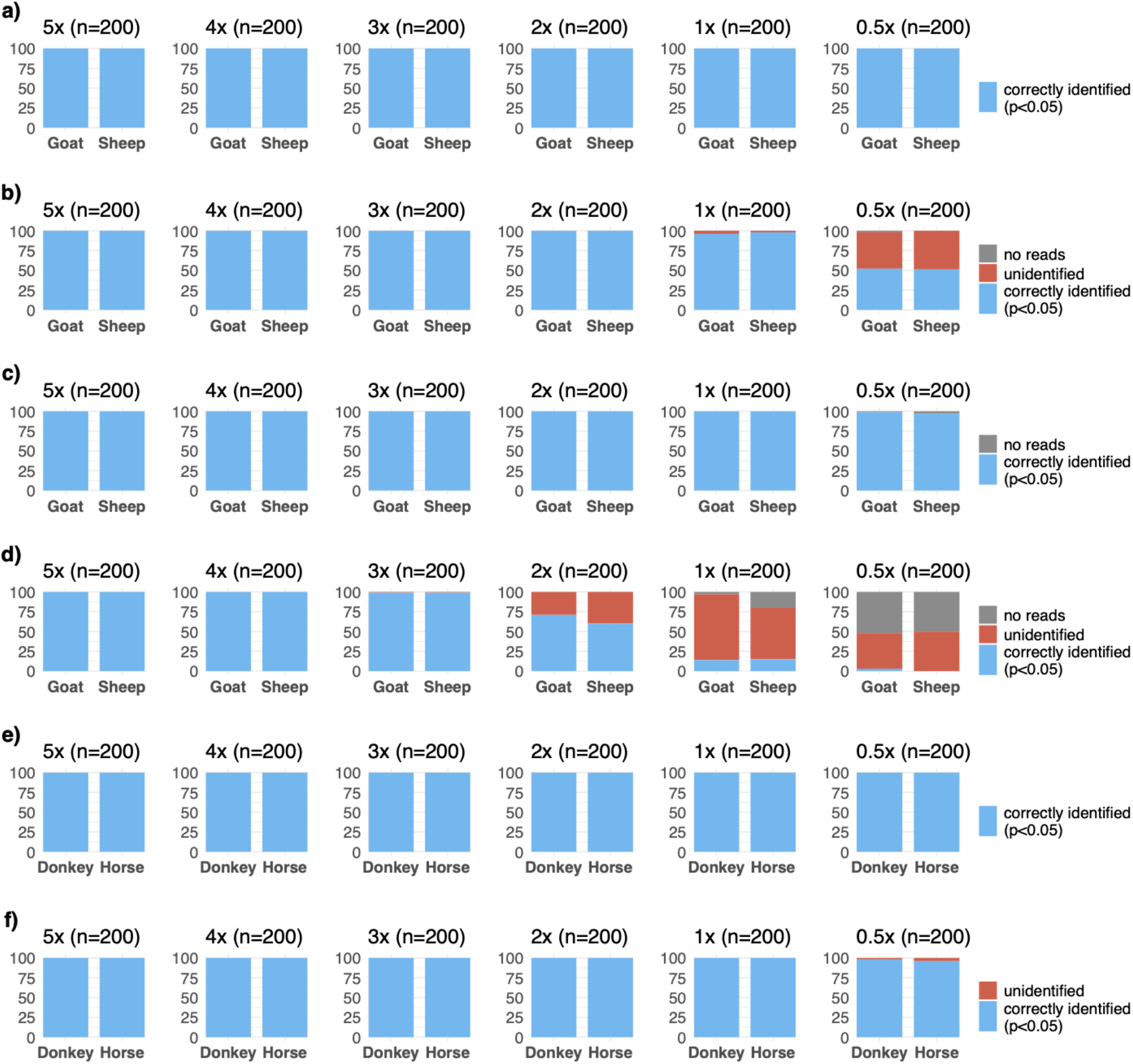
Results of the method applied to simulated ancient genomes at different coverages. Binomial test p-values for comparing the proportions of reads assigned to (a-d) sheep versus goat, and (e,f) and horse versus donkey. For sheep and goat, results are based on transversion substitutions without (a,b) and with (c,d) the exclusion of polymorphic sites. Results for both pairs are obtained through default (a,c,e) and “shared reads” approaches (b,d,f). n refers to the number of simulated genomes in each case (100 for each species in a pair). The height of the blue bar represents the number of simulated goat/donkey and sheep/horse genomes identified correctly with p<0.05, and the height of the red bar represents the number of unidentified cases. No cases were misidentified. The height of the grey bar represents the trials that did not contain any reads aligned to target sites, and thus could not be evaluated.

We also tested the performance of two more conservative approaches. First, we tried the “shared reads” option, which uses only a subset of reads aligned to both genomes; here the recall was >50% at 0.5x, but reached >95% at 1x coverage (Figure 3b). This low recall appears to be caused by the lack of power to reject the null hypothesis due to the majority reads being eliminated by the “shared reads” approach.

Second, we repeated the analysis after eliminating polymorphic sites from the target substitution set (target sites type 2 with n=120 positions), and obtained 98% precision at 0.5x coverage (Figure 3c). Through the “shared” reads approach and eliminating polymorphic sites (target sites type 2), we could assign taxa with 99% precision only at 3x coverage (Figure 3d). This was again apparently caused by reduced power due to using fewer sites and fewer reads.

We also performed the same analysis for n=1200 datasets of horse or donkey. Using n=117 transversion substitutions (target sites type 1), we again achieved 100% precision and recall in taxonomic assignment (Figure 3e). Using the “shared reads” option, we obtained 97% recall at 0.5x, and 100% recall at >0.5x coverage (Figure 3f). Thus, the performance of MTaxi was even more accurate for distinguishing horse and donkey, relative to distinguishing sheep and goat, despite the smaller target substitution set (see Discussion).

Overall, the simulations suggest that MTaxi can achieve full accuracy even at mtDNA coverages ≥0.5x. Conversely, limiting the analysis to subsets of reads aligned to both genomes or to non-polymorphic substitution positions reduces power, and does not increase accuracy.

### Application to samples of known species identity

#### Sheep and goat

We tested MTaxi on n=9 published ancient sheep (*Ovis aries* / *Ovis orientalis*) samples with mtDNA coverages >0.1x, and n=9 published ancient goat (*Capra hircus / Capra aegagrus*) samples with mtDNA coverages >0.3x (Table 2). The samples were produced in three different laboratories and varied in their mtDNA coverage. MTaxi yielded 100% accuracy for all 18 samples using the default approach (“all reads”) (Table 2). The probability of correct assignment by chance across all 18 MTaxi-classified specimens would be only 0.0003%, indicating the overall accuracy of our method.

**Table 2.** MTaxi results on sheep/goat genome data of known species identity using target sites type 1 with the default approach (“all reads”) The analysis was performed with n=197 transversion substitutions between sheep and goat, without excluding polymorphic sites. “Taxon” stands for known identity based on full genome data of the same sample (Table 1); “mtDNA coverage” shows coverage when mapping reads to the mtDNA reference of the original species (e.g. mtDNA coverage using the sheep reference for sheep data) after the duplicates have been removed; “Total assigned reads” refers to reads that could be mapped to both mtDNAs and the ones that could map only to one of the species’ references with high quality, overlapped target sites, and could be assigned to either species; “Sheep reads” and “Goat reads” show the number of reads that could be unambiguously assigned to either species; “p-value” shows the two-sided binomial test p-value for the proportion of sheep and goat reads being equal, and “Identified taxon” shows the final taxon assignment.

| Samples | Taxon | mtDNA coverage | Total reads | Sheep reads | Goat reads | p value | Identified taxon |
| --- | --- | --- | --- | --- | --- | --- | --- |
| TEP03 | Sheep | 12.5903 | 2643 | 1949 | 694 | <0.001 | Sheep |
| TEP62 | Sheep | 10.4074 | 3181 | 2555 | 626 | <0.001 | Sheep |
| TEP83 | Sheep | 5.0523 | 1474 | 1236 | 238 | <0.001 | Sheep |
| OBI014<br>(OB20-06) | Sheep | 4.29393 | 1043 | 1037 | 6 | <0.001 | Sheep |
| OBI018<br>(OB20-04) | Sheep | 1.12975 | 235 | 232 | 3 | <0.001 | Sheep |
| OBI013<br>(OB20-01) | Sheep | 0.578539 | 110 | 108 | 2 | <0.001 | Sheep |
| ULU26 | Sheep | 0.55579 | 178 | 171 | 7 | <0.001 | Sheep |
| ULU31 | Sheep | 0.428021 | 112 | 107 | 5 | <0.001 | Sheep |
| OBI017<br>(OB21-06) | Sheep | 0.100626 | 29 | 29 | 0 | <0.001 | Sheep |
| Acem1 | Goat | 10.9231 | 7969 | 751 | 7218 | <0.001 | Goat |
| AP45 | Goat | 0.267634 | 217 | 0 | 217 | <0.001 | Goat |
| Azer3 | Goat | 7.71564 | 5351 | 144 | 5207 | <0.001 | Goat |
| Direkli1 | Goat | 6.80549 | 5579 | 488 | 5091 | <0.001 | Goat |
| Direkli6 | Goat | 10.2286 | 10536 | 928 | 9608 | <0.001 | Goat |
| Gilat10 | Goat | 1.71545 | 171 | 1 | 170 | <0.001 | Goat |
| Shiqmim9 | Goat | 0.799651 | 84 | 1 | 83 | <0.001 | Goat |
| Kov27 | Goat | 2.99934 | 325 | 5 | 320 | <0.001 | Goat |
| Uiv17 | Goat | 0.909304 | 98 | 1 | 97 | <0.001 | Goat |

As observed in the simulations, using the “shared reads’’ approach did not improve accuracy, and we could correctly assign only 15 samples, while 3 samples with the lowest coverage had too few reads for assignment at p<0.05 (Table S1). One sheep sample (ULU31), with mtDNA coverage at 0.4x, had no reads overlapping the target sites, and thus could not be analysed at all. Interestingly, we observed 1-26% of reads misassigned with the default (“all reads”) approach. These could represent homoplasy, shared polymorphism, or PCR/sequencing error. However, they do not influence the final outcome.

Since the positions that remain polymorphic within species can introduce noise in downstream analyses, for sheep and goat, we also studied the performance of the method using target sites type 2 (excluding polymorphisms; n=120 sites). Using the default approach (“all reads”), this yielded 100% accuracy for all samples except one sheep, which had a coverage lower than 0.2x (Table S2). With the “shared reads” approach, 100% accuracy was achieved for only n=4 sheep and n=8 goat samples (Table S3). Meanwhile, one goat and three sheep samples with mtDNA coverage lower than 0.6x did not have any reads aligned to sheep and goat references that contained the target sites. We also noted that species-misassigned reads identified in the samples were not eliminated by this procedure (Tables 2-5). This result resonates with the above result from simulations, that removing polymorphic sites lowers statistical power but does not improve accuracy, at least in the case of sheep/goat assignment.

#### Horse and donkey

Applying MTaxi on n=5 horse and n=5 donkey samples, our method yielded 100% accuracy with both approaches (Table 3,S4). The overall rate of correct assignment in this sample set appears significant (one-sided binomial test p=0.001).

**Table 3.**
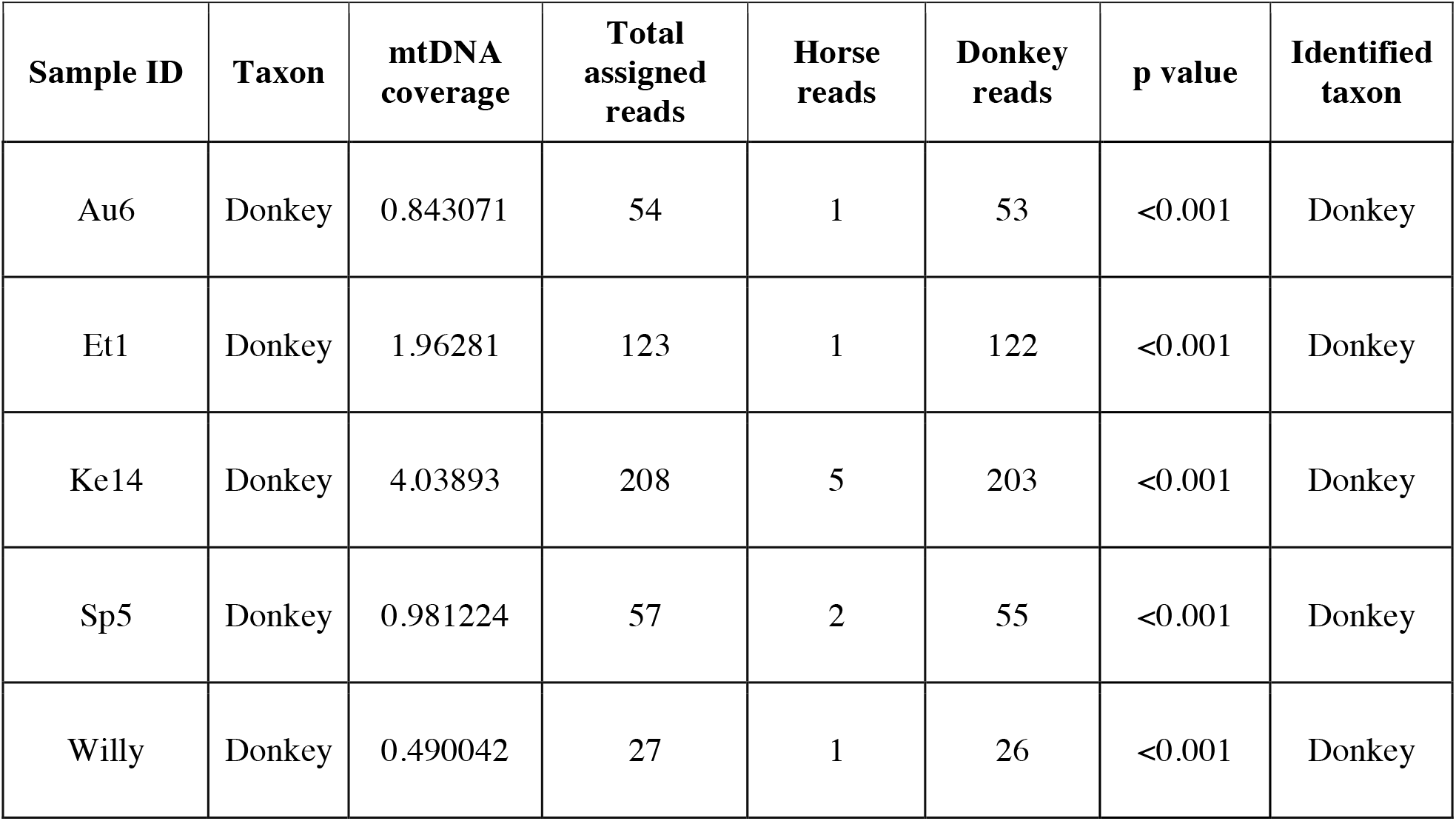
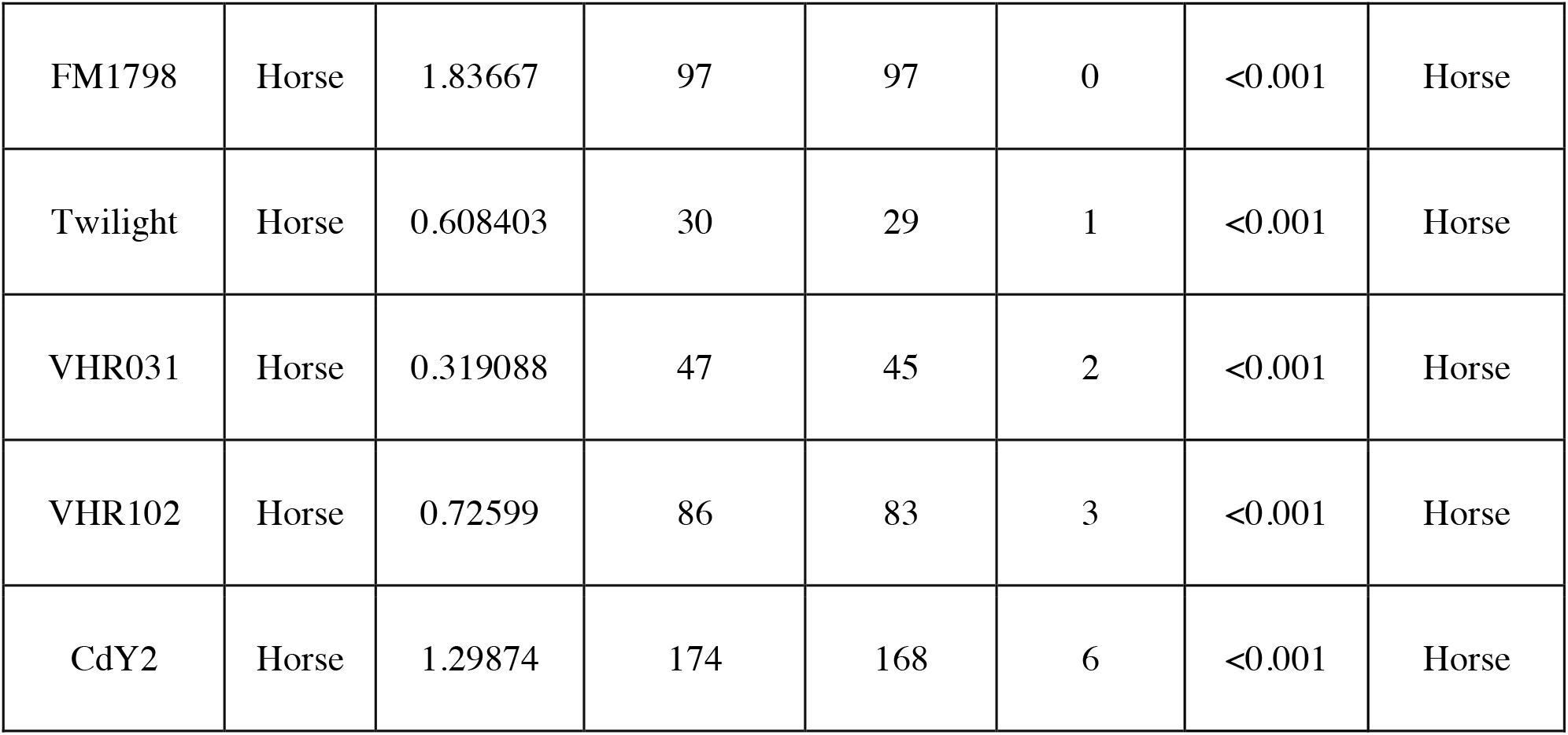
MTaxi results on horse/donkey genome data of known species identity using target sites type 1 with the default approach (“all reads”) The analysis was performed with n=117 transversion substitutions between horse and donkey (target sites type 1). “Taxon” stands for known identity based on full genome data of the same sample (Table 1); “mtDNA coverage” shows coverage when mapping reads to the mtDNA reference of the original species (e.g. mtDNA coverage using the horse reference for horse data) after the duplicates have been removed; “Total assigned reads” refers to reads that could be mapped to both mtDNAs and the ones that could map only to one of the species’ references with high quality, overlapped target sites, and could be assigned to either species; “Horse reads” and “Donkey reads” show the number of reads that could be unambiguously assigned to either species; “p-value” shows the two-sided binomial test p-value for the proportion of horse and donkey reads being equal, and “Identified taxon” shows the final taxon assignment.

### Whole genome comparative alignment

Comparative alignment can theoretically be a simple alternative to MTaxi when nuclear reference genomes are available. Here we explored the performance of comparative alignment using sheep/goat assignment as a model.

First, we observed that among ancient sheep BAM files used in this study, mapping results revealed inconsistencies in terms of the total number of bases aligning to each reference genome (Figure 4). Out of 9 sheep datasets with known species identity, only 4 showed a higher number of bases aligning to the sheep reference relative to the goat reference. However, we did not observe a similar inconsistency for the ancient goat samples, all of which had a higher number of bases mapped to the goat reference genome, most likely due to higher assembly quality of the goat reference. Unsurprisingly, the number of bases aligned to the nuclear genomes may not be an appropriate statistic for taxon identification between closely related taxon pairs (see Discussion).

**Figure 4.**
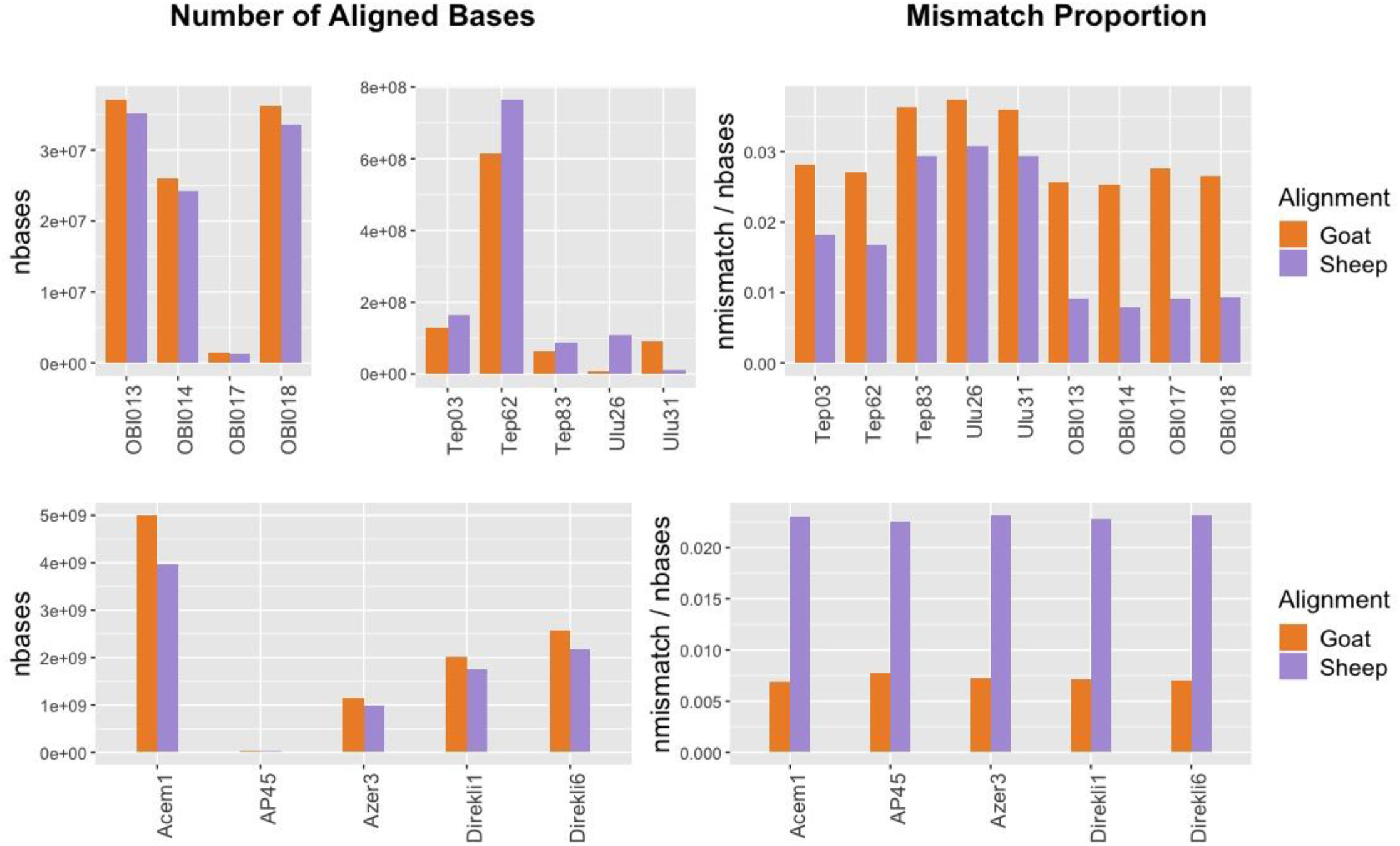
Total number of bases aligned to sheep versus goat reference genomes (left panels) and the mismatch proportions (right panels) for whole genome ancient sheep (upper panels) and goat (lower panels) samples.

We then analysed mismatch proportions of reads aligned to either genome. In all n=10 sheep and n=5 goat samples, including the lowest coverage samples with 0.0004x nuclear coverage, we found lower proportions of mismatches to their own reference genome. Again, this result is unlikely to happen by chance (one-sided binomial test p=3e-05). We note, however, that 3 of the sheep samples (TEP83, ULU26, ULU31) differ only marginally (by ∼0.7%) in their sheep vs. goat mismatch proportions. This suggests that comparative alignment can be an alternative to mitochondrial analysis in species identification, although our results imply that its success may not be guaranteed in all circumstances.

## Discussion

We showcase a simple but elaborate method to distinguish between closely related taxa using low coverage ancient DNA data, utilising mtDNA substitutions. Focusing on mtDNA is advantageous both in terms of high copy number and of greater variability (see Introduction). Also, the mtDNA/nuclear DNA ratio has been stated to correlate positively with a decrease in endogenous DNA content (22), suggesting that it should be more likely to obtain higher amounts of mtDNA than nuclear DNA in particularly poorly preserved samples.

Simulation results showed that MTaxi can distinguish sheep vs. goat with full accuracy at mtDNA coverages ≥0.5x. We also obtained 100% correct results with 18 ancient samples of known identity, which had mtDNA coverages between 0.1x-12x (one sided binomial tests p = 3e-06). Likewise, simulations of ancient horse and donkey data yielded 100% accurate results at mtDNA coverages ≥0.5x, while downsampled modern and ancient domestic equid samples (n=10) of known species identity were also assigned fully correctly (one-sided binomial test p=0.001). Overall, MTaxi appears as a simple and efficient tool for correct taxon identification using ultra-low coverage shotgun sequencing data.

Meanwhile, our results suggested that conservative modifications of the pipeline that involve limiting the analysis to “shared reads” or excluding polymorphic sites did not improve performance, but on the contrary reduced statistical power and recall.

We note that MTaxi is successful even at mitochondrial coverages of 0.5x, which is a level frequently reached in low-coverage sequencing experiments when there exists 1% endogenous DNA. For example, in the aforementioned shotgun sequencing dataset (see Introduction), we had n=226 ancient mammalian samples with 1-10% endogenous DNA (median = 3%), and each library sequenced to a size of up to 50 million total reads per sample (median = 415,146 reads); within this set 50% of the libraries reached mitochondrial DNA coverages ≥0.5x, sufficient for effective identification by MTaxi (while 62% and 84% of the libraries reached ≥0.3x and ≥0.1x, respectively).

Our observations on comparative alignment were also notable. The comparison of total number of bases mapped to sheep and goat reference genomes showed that mapping frequencies can be deceiving, even when analysing whole genome data. A sheep FASTQ file can align more widely to the goat reference genome, and the degree to which this occurs seems to vary among samples. Meanwhile, all the goat samples had higher numbers of bases mapped to the goat reference genome. The reason for the observed differences between the performance of goat and sheep samples in alignment of their respective genomes could be related to variability in reference genome qualities and/or polymorphism between the species (indeed, the N50 of the goat genome ARS1 is 26,244,591 while that for the sheep genome Oar_v3.1 is 40,376). More generally, this result indicates that taxon identification using only the number of aligned bases in comparative alignment is not reliable.

The comparison of mismatch proportions in comparative alignment, on the other hand, appears to be a relatively robust approach based on our empirical sample of 15 sheep and goat samples, even at a nuclear coverage of 0.0004x (also a value typically displayed in low coverage sequencing experiments). This could be a simple solution for taxon identification if reference genomes are available for both taxa. Still, our observation that mismatch proportions can vary only marginally in some sheep samples mapped to goat (e.g. TEP83 and ULU26 in Fig. 5), calls for caution in using this strategy.

MTaxi would be expected to perform on any species pairs with a degree of divergence comparable to that of sheep and goat, and would be particularly convenient when reference nuclear genomes of one of the species is lacking, which precludes comparative alignment. Candidate taxa that pose challenges for zooarchaeological identification include several mammal species in families Cervidae (deer), Leporidae (rabbit/hare) and Bovidae (cattle/bison), and birds (45-47). Horse and donkey are another such pair, on which we checked the performance of our method. Compared to the Zonkey pipeline (21), designed to classify ancient equid samples, MTaxi does not require a reference panel and is solely based on mitochondrial DNA data, hence an easier and faster method of classification.

In summary, the performance of MTaxi will depend on various factors, including evolutionary divergence and reference genome qualities of the species pairs, but we expect it to be an effective tool in various settings, as long as mitochondrial introgression can be excluded. We also note that its parameters and the data processing steps can be fine tuned to adjust for particularities of the species in question, such as the exclusion of polymorphic sites.

## Data Availability

All data underlying the results are available as part of the article and no additional source data are required.

## Software Availability

Source code is available from https://github.com/goztag/MTaxi

## Acknowledgements

We thank all the members of METU CompEvo group for their helpful suggestions.

## References

1. Braje TJ, Rick TC, Szpak P, Newsome SD, McCain JM, Elliott Smith EA, et al. Historical ecology and the conservation of large, hermaphroditic fishes in Pacific coast kelp forest ecosystems. Science Advances. 2017;3(2).

2. Costa T, Barri F. Lama guanicoe remains from the Chaco ecoregion (Córdoba, Argentina): An osteological approach to the characterization of a relict wild population. PLOS ONE. 2018;13(4).

3. Murray MS. Zooarchaeology and Arctic Marine Mammal Biogeography, conservation, and Management. Ecological Applications. 2008 Mar 18;18(sp2).

4. Gifford-Gonzalez D. Zooarchaeology and Ecology: Mortality Profiles, Species Abundance, Diversity. In: An introduction to zooarchaeology. Cham: Springer; 2018. p. 475–501.

5. Wolverton S, Lyman RL. Applied Zooarchaeology History, Value, and Use. In: Conservation Biology and Applied Zooarchaeology. Tucson: University of Arizona Press; 2012.

6. Steele TE. The contributions of animal bones from archaeological sites: The past and future of zooarchaeology. Journal of Archaeological Science. 2015;56:168–76.

7. LeFebvre MJ, Sharpe AE. Contemporary challenges in zooarchaeological specimen identification. Zooarchaeology in Practice. 2017;:35–57.

8. Wolfhagen J, Price MD. A probabilistic model for distinguishing between sheep and goat postcranial remains. Journal of Archaeological Science: Reports. 2017;12:625–31.

9. Parson W, Pegoraro K, Niedersttter H, Fger M, Steinlechner M. Species identification by means of the cytochrome b gene. International Journal of Legal Medicine. 2000;114(1-2):23–8.

10. Newman ME, Parboosingh JS, Bridge PJ, Ceri H. Identification of archaeological animal bone by PCR/DNA analysis. Journal of Archaeological Science. 2002;29(1):77–84.

11. Kahila Bar-Gal G, Ducos P, Kolska Horwitz L. The application of ancient DNA analysis to identify neolithic Caprinae: A case study from the site of Hatoula, Israel. International Journal of Osteoarchaeology. 2003;13(3):120–31.

12. Balasse M, Ambrose SH. Distinguishing sheep and goats using dental morphology and stable carbon isotopes in C4 Grassland Environments. Journal of Archaeological Science. 2005;32(5):691–702.

13. Buckley M, Kansa SW. Collagen fingerprinting of archaeological bone and teeth remains from Domuztepe, south eastern Turkey. Archaeological and Anthropological Sciences. 2011;3(3):271–80.

14. Lan T, Lin Y, Njaramba-Ngatia J, Guo X, Li R, Li H, et al. Improving species identification of ancient mammals based on next-generation sequencing data. Genes. 2019;10(7):509.

15. Orlando L, Allaby R, Skoglund P, Der Sarkissian C, Stockhammer PW, Ávila-Arcos MC, et al. Ancient DNA analysis. Nature Reviews Methods Primers. 2021;1(1).

16. Allentoft ME, Collins M, Harker D, Haile J, Oskam CL, Hale ML, et al. The half-life of DNA in bone: Measuring decay kinetics in 158 dated fossils. Proceedings of the Royal Society B: Biological Sciences. 2012;279(1748):4724–33.

17. Ottoni C, Koon HE, Collins MJ, Penkman KE, Rickards O, Craig OE. Preservation of ancient DNA in thermally damaged archaeological bone. Naturwissenschaften. 2008;96(2):267–78.

18. Carpenter ML, Buenrostro JD, Valdiosera C, Schroeder H, Allentoft ME, Sikora M, et al. Pulling out the 1%: Whole-genome capture for the targeted enrichment of ancient DNA sequencing libraries. The American Journal of Human Genetics. 2013;93(5):852–64.

19. Yurtman E, Özer O, Yüncü E, Dagtas ND, Koptekin D, Çakan YG, et al. Archaeogenetic analysis of neolithic sheep from Anatolia suggests a complex demographic history since domestication. Communications Biology. 2021;4(1).

20. Taylor WT, Pruvost M, Posth C, Rendu W, Krajcarz MT, Abdykanova A, et al. Evidence for early dispersal of domestic sheep into Central Asia. Nature Human Behaviour. 2021;5(9):1169–79.

21. Schubert M, Mashkour M, Gaunitz C, Fages A, Seguin-Orlando A, Sheikhi S, et al. Zonkey: A simple, accurate and sensitive pipeline to genetically identify equine F1-hybrids in archaeological assemblages. Journal of Archaeological Science. 2017;78:147–57.

22. Furtwängler A, Reiter E, Neumann GU, Siebke I, Steuri N, Hafner A, et al. Ratio of mitochondrial to nuclear DNA affects contamination estimates in ancient DNA analysis. Scientific Reports. 2018;8(1).

23. Brown WM, George M, Wilson AC. Rapid evolution of animal mitochondrial DNA. Proceedings of the National Academy of Sciences. 1979;76(4):1967–71.

24. NCBI Genome Data viewer [Internet]. National Center for Biotechnology Information. U.S. National Library of Medicine; [cited 2021 Dec 24]. Available from: https://www.ncbi.nlm.nih.gov/genome/gdv/?org=homo-sapiens&group=bilateria

25. Genome list - genome - NCBI [Internet]. National Center for Biotechnology Information. U.S. National Library of Medicine; [cited 2021Dec24]. Available from: https://www.ncbi.nlm.nih.gov/genome/browse#!/organelles/

26. Daly KG, Delser PM, Mullin VE, Scheu A, Mattiangeli V, Teasdale MD, et al. Ancient goat genomes reveal mosaic domestication in the Fertile Crescent. Science. 2018;361(6397):85–8.

27. Fages A, Hanghøj K, Khan N, Gaunitz C, Seguin-Orlando A, Leonardi M, et al. Tracking five millennia of horse management with extensive ancient genome time series. Cell. 2019;177(6).

28. Orlando L, Ginolhac A, Zhang G, Froese D, Albrechtsen A, Stiller M, et al. Recalibrating equus evolution using the genome sequence of an early middle pleistocene horse. Nature. 2013;499(7456):74–8.

29. Der Sarkissian C, Ermini L, Schubert M, Yang MA, Librado P, Fumagalli M, et al. Evolutionary genomics and conservation of the endangered przewalski’s horse. Current Biology. 2015;25(19):2577–83.

30. Renaud G, Petersen B, Seguin-Orlando A, Bertelsen MF, Waller A, Newton R, et al. Improved de novo genomic assembly for the domestic donkey. Science Advances. 2018;4(4).

31. Wang C, Li H, Guo Y, Huang J, Sun Y, Min J, et al. Donkey genomes provide new insights into domestication and selection for coat color. Nature Communications. 2020;11(1).

32. H. Pagès, P. Aboyoun, R. Gentleman and S. DebRoy. Biostrings:Efficient manipulation of biological strings. R package version 2.54.0. 2019.

33. Broughton RE, Stanley SE, Durrett RT. Quantification of homoplasy for nucleotide transitions and transversions and a reexamination of assumptions in weighted phylogenetic analysis. Systematic Biology. 2000;49(4):617–27.

34. Page AJ, Taylor B, Delaney AJ, Soares J, Seemann T, Keane JA, et al. SNP-sites: rapid efficient extraction of SNPs from multi-FASTA alignments. Microbial Genomics. 2016;2(4).

35. Shi N-N, Fan L, Yao Y-G, Peng M-S, Zhang Y-P. Mitochondrial genomes of domestic animals need scrutiny. Molecular Ecology. 2014;23(22):5393–7.

36. Grant JR, Stothard P. The CGView Server: a comparative genomics tool for circular genomes. Nucleic Acids Research. 2008;36(Web Server).

37. Schubert M, Lindgreen S, Orlando L. Adapterremoval v2: Rapid adapter trimming, identification, and read merging. BMC Research Notes. 2016;9(1).

38. Li H, Durbin R. Fast and accurate long-read alignment with Burrows–Wheeler transform. Bioinformatics. 2010;26(5):589–95.

39. Kircher M. Analysis of high-throughput ancient DNA sequencing data. Methods in Molecular Biology. 2011;:197–228.

40. Li H, Handsaker B, Wysoker A, Fennell T, Ruan J, Homer N, et al. The sequence alignment/map format and SAMtools. Bioinformatics. 2009;25(16):2078–9.

41. Skoglund P, Malmström H, Omrak A, Raghavan M, Valdiosera C, Günther T, et al. Genomic diversity and admixture differs for stone-age Scandinavian foragers and Farmers. Science. 2014;344(6185):747–50.

42. Quinlan AR, Hall IM. BEDTools: a flexible suite of utilities for comparing genomic features. Bioinformatics. 2010;26(6):841–2.

43. Dale RK, Pedersen BS, Quinlan AR. Pybedtools: A flexible python library for manipulating genomic datasets and Annotations. Bioinformatics. 2011;27(24):3423–4.

44. Renaud G, Hanghøj Kx, Willerslev E, Orlando L. gargammel: a sequence simulator for ancient DNA. Bioinformatics. 2016;

45. Wolverton S. Data quality in zooarchaeological faunal identification. Journal of Archaeological Method and Theory. 2013;20:381–96.

46. Yang DY, Cannon A, Saunders SR. DNA species identification of archaeological salmon bone from the Pacific Northwest Coast of North America. Journal of Archaeological Science. 2004;31:619–31.

47. Bochenski ZM. Identification of skeletal remains of closely related species: the pitfalls and solutions. Journal of Archaeological Science. 2008;35:1247–50.

