## Supplemental Tables for "MTaxi : A comparative tool for taxon identification of ultra low coverage ancient genomes"

**Table S1. MTaxi results on sheep/goat genome data of known species identity using target sites type 1 using the "shared reads" approach**

The analysis was performed with n=197 transversion substitutions between sheep and goat, without excluding polymorphic sites. "Taxon" stands for known identity based on full genome data of the same sample (Table 1); "mtDNA coverage" shows coverage when mapping reads to the mtDNA reference of the original species (e.g. mtDNA coverage using the sheep reference for sheep data) after the duplicates have been removed; "Total assigned reads" refers to reads that could be mapped to both mtDNA with high quality, overlapped target sites, and could be assigned to either species; "Sheep reads" and "Goat reads" show the number of reads that could be unambiguously assigned to either species; "p-value" shows the two-sided binomial test p-value for the proportion of sheep and goat reads being equal, and "Identified taxon" shows the final taxon assignment. (□) indicates that the sample that did not contain any reads aligned to target sites, and thus could not be evaluated.

| Sample ID | Taxon | mtDNA coverage | Total assigned reads | Sheep reads | Goat reads | p-value | Identified taxon |
|---|---|---|---|---|---|---|---|
| TEP03 | Sheep | 12.5903 | 70 | 70 | 0 | <0.001 | Sheep |
| TEP62 | Sheep | 10.4074 | 52 | 52 | 0 | <0.001 | Sheep |
| TEP83 | Sheep | 5.0523 | 42 | 42 | 0 | <0.001 | Sheep |
| OBI014 (OB20-06) | Sheep | 4.29393 | 30 | 30 | 0 | <0.001 | Sheep |
| OBI018 (OB20-04) | Sheep | 1.12975 | 12 | 12 | 0 | <0.001 | Sheep |
| OBI013 (OB20-01) | Sheep | 0.578539 | 5 | 5 | 0 | 0.0625 | - |
| ULU26 | Sheep | 0.55579 | 6 | 6 | 0 | 0.03125 | Sheep |
| ULU31□ | Sheep | 0.428021 | 0 | 0 | 0 | NA | - |
| OBI017 (OB21-06) | Sheep | 0.100626 | 1 | 1 | 0 | 1.0 | - |
| Acem1 | Goat | 10.9231 | 255 | 0 | 255 | <0.001 | Goat |

| | | | | | | | |
|---|---|---|---|---|---|---|---|
| AP45 | Goat | 0.267634 | 8 | 0 | 8 | 0.00781 | Goat |
| Azer3 | Goat | 7.71564 | 246 | 0 | 246 | <0.001 | Goat |
| Direkli1 | Goat | 6.80549 | 215 | 13 | 202 | <0.001 | Goat |
| Direkli6 | Goat | 10.2286 | 363 | 15 | 348 | <0.001 | Goat |
| Gilat10 | Goat | 1.71545 | 32 | 0 | 32 | <0.001 | Goat |
| Shiqmim9 | Goat | 0.799651 | 14 | 0 | 14 | <0.001 | Goat |
| Kov27 | Goat | 2.99934 | 62 | 0 | 62 | <0.001 | Goat |
| Uiv17 | Goat | 0.909304 | 19 | 0 | 19 | <0.001 | Goat |

**Table S2. MTaxi results on sheep/goat genome data of known species identity using target sites type 2 with the default approach ("all reads")**

The analysis was performed with n=120 transversion substitutions (target sites type 2) between sheep and goat, excluding polymorphic sites. "Taxon" stands for known identity based on full genome data of the same sample (Table 1); "mtDNA coverage" shows coverage when mapping reads to the mtDNA reference of the original species (e.g. mtDNA coverage using the sheep reference for sheep data) after the duplicates have been removed; "Total assigned reads" refers to reads that could be mapped to both mtDNAs and the ones that could map only to one of the species' references with high quality, overlapped target sites, and could be assigned to either species; "Sheep reads" and "Goat reads" show the number of reads that could be unambiguously assigned to either species; "p-value" shows the two-sided binomial test p-value for the proportion of sheep and goat reads being equal, and "Identified taxon" shows the final taxon assignment. (□) indicates that the sample that did not contain any reads aligned to target sites, and thus could not be evaluated.

| Sample ID | Taxon | mtDNA coverage | Total assigned reads | Sheep reads | Goat reads | p value | Identified taxon |
|---|---|---|---|---|---|---|---|

| | | | | | | | |
|---|---|---|---|---|---|---|---|
| TEP03 | Sheep | 12.5903 | 2291 | 1659 | 632 | <0.001 | Sheep |
| TEP62 | Sheep | 10.4074 | 2735 | 2195 | 540 | <0.001 | Sheep |
| TEP83 | Sheep | 5.0523 | 1320 | 1066 | 254 | <0.001 | Sheep |
| OBI014 (OB20-06) | Sheep | 4.29393 | 837 | 835 | 2 | <0.001 | Sheep |
| OBI018 (OB20-04) | Sheep | 1.12975 | 186 | 184 | 2 | <0.001 | Sheep |
| OBI013 (OB20-01) | Sheep | 0.578539 | 85 | 85 | 0 | <0.001 | Sheep |
| ULU26 | Sheep | 0.55579 | 124 | 115 | 9 | <0.001 | Sheep |
| ULU31 | Sheep | 0.428021 | 89 | 89 | 0 | <0.001 | Sheep |
| OBI017□ (OB21-06) | Sheep | 0.100626 | 0 | 0 | 0 | NA | - |
| Acem1 | Goat | 10.9231 | 6627 | 845 | 5782 | <0.001 | Goat |
| AP45 | Goat | 0.267634 | 177 | 0 | 177 | <0.001 | Goat |
| Azer3 | Goat | 7.71564 | 4188 | 196 | 3992 | <0.001 | Goat |
| Direkli1 | Goat | 6.80549 | 4568 | 530 | 4038 | <0.001 | Goat |
| Direkli6 | Goat | 10.2286 | 8545 | 995 | 7550 | <0.001 | Goat |
| Gilat10 | Goat | 1.71545 | 136 | 1 | 135 | <0.001 | Goat |

| | | | | | | | |
|---|---|---|---|---|---|---|---|
| Shiqmim9 | Goat | 0.799651 | 70 | 1 | 69 | <0.001 | Goat |
| Kov27 | Goat | 2.99934 | 251 | 5 | 246 | <0.001 | Goat |
| Uiv17 | Goat | 0.909304 | 87 | 1 | 86 | <0.001 | Goat |

**Table S3. MTaxi results on sheep/goat genome data of known species identity using target sites type 2 with the "shared reads" approach**

The analysis was performed with n=120 transversion substitutions (target sites type 2) between sheep and goat, excluding polymorphic sites. "Taxon" stands for known identity based on full genome data of the same sample (Table 1); "mtDNA coverage" shows coverage when mapping reads to the mtDNA reference of the original species (e.g. mtDNA coverage using the sheep reference for sheep data) after the duplicates have been removed; "Total assigned reads" refers to reads that could be mapped to both mtDNA with high quality, overlapped target sites, and could be assigned to either species; "Sheep reads" and "Goat reads" show the number of reads that could be unambiguously assigned to either species; "p-value" shows the two-sided binomial test p-value for the proportion of sheep and goat reads being equal, and "Identified taxon" shows the final taxon assignment. (☐) indicates that the sample that did not contain any reads aligned to target sites, and thus could not be evaluated.

| Sample ID | Taxon | mtDNA coverage | Total assigned reads | Sheep reads | Goat reads | p value | Identified taxon |
|---|---|---|---|---|---|---|---|
| TEP03 | Sheep | 12.5903 | 34 | 34 | 0 | <0.001 | Sheep |
| TEP62 | Sheep | 10.4074 | 36 | 36 | 0 | <0.001 | Sheep |
| TEP83 | Sheep | 5.0523 | 14 | 14 | 0 | <0.001 | Sheep |
| OBI014 (OB20-06) | Sheep | 4.29393 | 14 | 14 | 0 | <0.001 | Sheep |

| | | | | | | | |
|---|---|---|---|---|---|---|---|
| OBI018 (OB20-04) | Sheep | 1.12975 | 1 | 1 | 0 | 1.0 | - |
| OBI013 (OB20-01) | Sheep | 0.578539 | 2 | 2 | 0 | 0.5 | - |
| ULU26□ | Sheep | 0.55579 | 0 | 0 | 0 | NA | - |
| ULU31□ | Sheep | 0.428021 | 0 | 0 | 0 | NA | - |
| OBI017 □ (OB21-06) | Sheep | 0.100626 | 0 | 0 | 0 | NA | - |
| Acem1 | Goat | 10.9231 | 122 | 0 | 122 | <0.001 | Goat |
| AP45 | Goat | 0.267634 | 3 | 0 | 3 | 0.25 | - |
| Azer3 | Goat | 7.71564 | 87 | 0 | 87 | <0.001 | Goat |
| Direkli1 | Goat | 6.80549 | 92 | 13 | 79 | <0.001 | Goat |
| Direkli6 | Goat | 10.2286 | 174 | 15 | 159 | <0.001 | Goat |
| Gilat10 | Goat | 1.71545 | 24 | 0 | 24 | <0.001 | Goat |
| Shiqmim9 | Goat | 0.799651 | 9 | 0 | 9 | 0.003906 | Goat |
| Kov27 | Goat | 2.99934 | 41 | 0 | 41 | <0.001 | Goat |
| Uiv17 | Goat | 0.909304 | 17 | 0 | 17 | <0.001 | Goat |

**Table S4. MTaxi results on horse/donkey genome data of known species identity using target sites type 1 with the "shared reads" approach**

The analysis was performed with n=117 transversion substitutions between horse and donkey (target sites type 1). "Taxon" stands for known identity based on full genome data of the same sample (Table 1); "mtDNA coverage" shows coverage when mapping reads to the mtDNA reference of the original species (e.g. mtDNA coverage using the horse reference for horse data) after the duplicates have been removed; "Total assigned reads" refers to reads that could be mapped to both mtDNA with high quality, overlapped target sites, and could be assigned to either species; "Horse reads" and "Donkey reads" show the number of reads that could be unambiguously assigned to either species; "p-value" shows the two-sided binomial test p-value for the proportion of horse and donkey reads being equal, and "Identified taxon" shows the final taxon assignment.

| Sample ID | Taxon | mtDNA coverage | Total assigned reads | Horse reads | Donkey reads | p value | Identified taxon |
|---|---|---|---|---|---|---|---|
| Au6 | Donkey | 0.843071 | 47 | 1 | 46 | <0.001 | Donkey |
| Et1 | Donkey | 1.96281 | 106 | 0 | 106 | <0.001 | Donkey |
| Ke14 | Donkey | 4.03893 | 186 | 4 | 182 | <0.001 | Donkey |
| Sp5 | Donkey | 0.981224 | 52 | 1 | 52 | <0.001 | Donkey |
| Willy | Donkey | 0.490042 | 20 | 1 | 19 | <0.001 | Donkey |
| FM1798 | Horse | 1.83667 | 85 | 85 | 0 | <0.001 | Horse |
| Twilight | Horse | 0.608403 | 21 | 20 | 1 | <0.001 | Horse |
| VHR031 | Horse | 0.319088 | 18 | 16 | 2 | 0.001312 | Horse |
| VHR102 | Horse | 0.72599 | 26 | 24 | 2 | <0.001 | Horse |

| | | | | | | | |
|---|---|---|---|---|---|---|---|
| CdY2 | Horse | 1.29874 | 40 | 40 | 0 | <0.001 | Horse |
